# N-terminal modification of Histone H3 inhibits H3K27M-mediated loss of H3K27 trimethylation

**DOI:** 10.1101/2025.01.24.634760

**Authors:** Nicolas Poux, Gabriel E Boyle, Daren F Zhang, Jacqueline Caplan, Adam F Kebede, Apichaya Sethaudom, Isabella Gainey, Olohireme E Famous, Jay Sarthy, Pratiti Bandopadhayay

## Abstract

The use of short epitope tags is widespread in the study of histone biology as they allow for the antibody-mediated detection and pulldown of post-translationally modified histones or exogenous histone transgenes. However, H3 is particularly sensitive to sequence modification and the addition of epitope tags may interfere with the native function of H3. Here, we use the known relationship between lysine-to-methionine K27M mutations and the loss of trimethylation at the H3 K27 residue to test whether the addition of epitope tags to the N or C terminus of H3 affects the levels of histone H3 post-translational modifications (PTMs). We find that all tested N-terminal tags abrogate the H3K27M-mediated loss of H3K27 trimethylation despite robust deposition in chromatin. These results suggest that the addition of epitope tags to the N-terminus of H3 should be performed with caution, and these findings may be of particular interest for the study of H3-driven cancers, like diffuse midline gliomas.

## Introduction

Post-translational modification of the N-terminal tail of histone H3 is a central mechanism driving the formation of epigenetic states^1^. Mutations in this tail sequence alter the ability of histone modifiers and readers to bind to the tail, leading to epigenetic dysregulation, and have been associated with developmental disorders and cancers. Lysine 27 is a key regulatory site and lysine to methionine (K-to-M) mutations at this position (H3K27M) are the initiating genomic event in diffuse midline gliomas (DMGs), a fatal pediatric brain tumor, and have been shown to block the deposition of the repressive H3K27 trimethylation mark (H3K27me3)^2–4^.

The sequence similarity of H3K27M to wild-type H3 renders the study of the mutation difficult without the use of epitope tags. These tags are short amino acid sequences that provide a specific binding motif for antibody-mediated detection and pulldown of proteins. Epitope tags have been foundational in establishing the oncogenic role of H3K27M right from the earliest cellular models demonstrating that fractional expression of H3K27M induced the loss of H3K27me3^4^, to more recent papers studying the mechanisms and extent of epigenetic reprogramming induced by H3K27M^5–7^.

However, the heavily conserved histone proteins show an exquisite sensitivity to sequence alterations. Even the five amino acid differences between the H3 variants H3.1 and H3.3 are sufficient for them to be deposited into nucleosomes using separate molecular machinery^8–11^. As such, it is possible that the addition of epitope tags may interfere with the function of endogenous H3 or the H3K27M mutant.

There is little consensus on the best practices for the use of epitope tags in the DMG literature. The most commonly used tags, FLAG, HA, and V5, possess different features that might uniquely affect H3 function: FLAG is enriched for negative side chains, HA contains many bulky tyrosines and prolines, and V5 is nearly twice as large as the other two. Additionally, the location of the tag carries different implications: the H3 N-terminus contains the unstructured histone tail where post-translational modifications are deposited, whereas the C-terminus forms the histone core that determines the strength of H3-DNA binding interactions. Many seminal papers in the DMG field differ in the identity and location of the tag used, introducing a potential source of variability across findings^4,7,12–14^. Since chromatin writer activity is regulated by subtle biochemical changes introduced by post-translational modifications (PTMs) at the H3 tail^6,15^, we reasoned that appending epitope tags to this same region may alter histone binding-protein activity and function.

We thus hypothesized that N-terminal epitope tags disrupt essential functions of the histone H3 tail. In this study, we use the established relationship between the H3K27M mutation and loss of H3K27me3 levels to test whether the addition of N-terminal epitope tags alters the effects of this mutation.

## Results

We sought to generate a platform to flexibly test different combinations of H3 variants and epitope tags (Figure 1). Many published DMG models harness lentivirus-mediated overexpression systems to introduce the mutant H3K27M into various human cell lines^4,12–14^, leveraging the dominant-negative effect of H3K27M to produce a reduction in H3K27me3 with fractional expression of the mutant histone. We first generated such models across several cell lines to express H3.3, including both the wild-type and K27M mutant transcripts, without any tag. We initially focused on induced pluripotent stem cell-derived human neural stem cells (hNSCs). In this isogenic setting, expression of untagged K27M mutant H3.3 (H3.3K27M) caused a significant reduction in H3K27me3 levels compared to untransduced cells (Figure 1a, 21.2% of untransduced control, p=0.003 across 3 independent replicates), whereas cells overexpressing the H3.3 wild-type control maintained similar H3K27me3 levels (129.7% of untransduced cells, p=0.246). The level of abundance of H3K27me3 in H3.3K27M expressing hNSCs was similar to that observed in patient-derived H3.3K27M mutant DMG cell lines BT245 and SF7761. We conclude that the overexpression of H3.3K27M is sufficient to suppress the polycomb repressive mark H3K27me3, as previously reported^4,7,12^.

**Figure 1.**
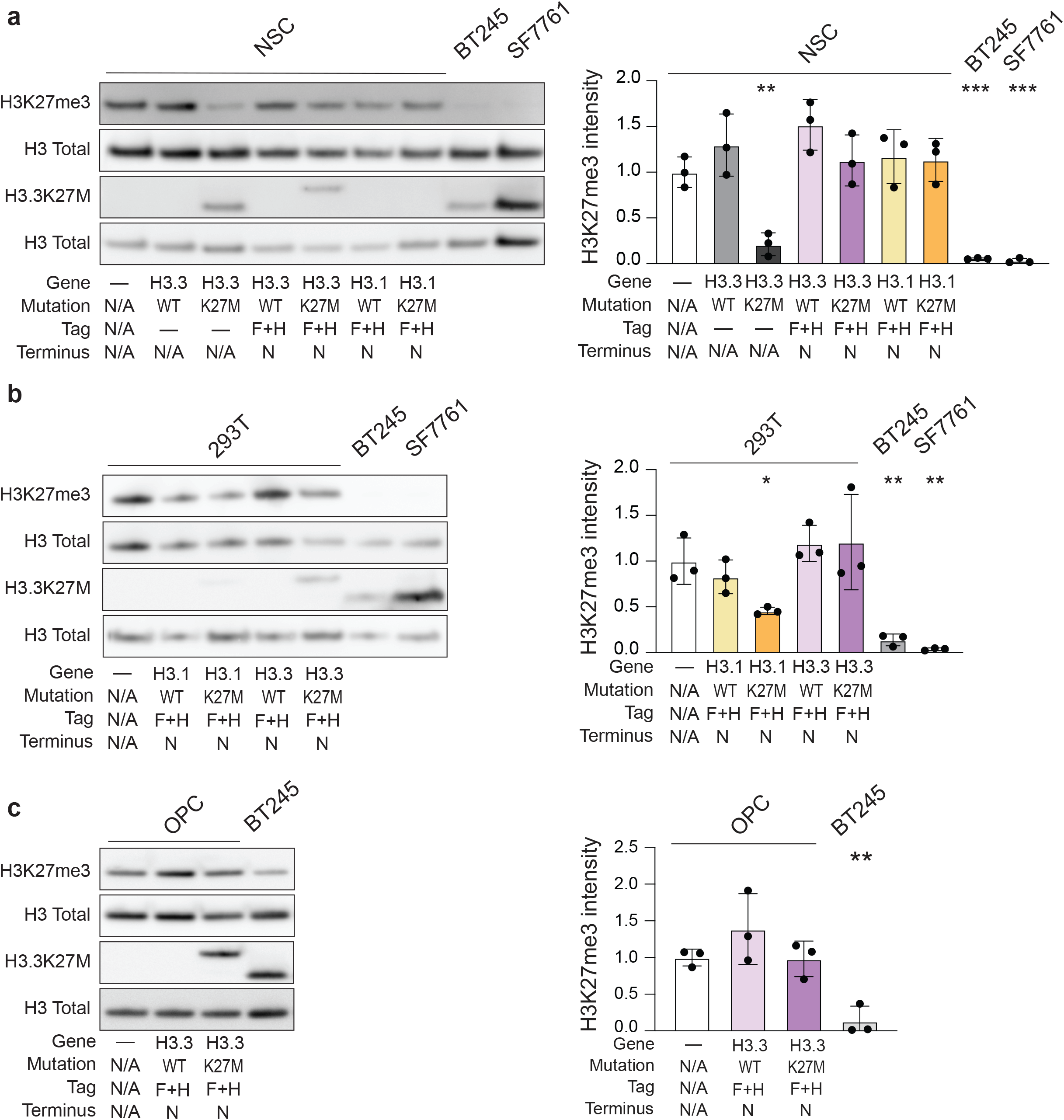
N-terminal FLAG+HA tag inhibits H3K27M-mediated reduction in H3K27me3 levels. **a.** Left: Western immunoblot of acid-extracted lysates from transduced hNSCs or the BT245 and SF7761 patient-derived DMG cell lines. Right: quantification of H3K27me3 : H3 ratios normalized to untransduced NSC control, n = 3 biological replicates. **b.** Left: Western immunoblot of acid-extracted lysates from transduced HEK293T cells compared to BT245 and SF7761. Right: quantification of H3K27me3 : H3 ratios normalized to untransduced HEK293T control, n = 3 biological replicates. **c.** Left: Western immunoblot of acid-extracted lysates from transduced hOPCs or BT245 cell lines. Right: quantification of H3K27me3 : H3 ratios normalized to untransduced hOPC control, n = 3 biological replicates. Significance testing performed with two-sided unpaired t-test against respective untransduced sample. Error bars, mean ± SEM of three replicates per condition. * = p<0.05, **=p<0.01, ***=p<0.001.

Our NSCs provided a model in which we could systematically evaluate the effects of epitope tags. We next evaluated whether similar suppression of H3K27me3 is observed with the introduction of N-terminal tags. We generated lentivirus overexpression constructs encoding either wild-type H3.1 or H3.3, or the K27M mutants (H3.1K27M and H3.3K27M). We then added a FLAG-HA tag, which was used in the original cellular models describing H3K27M-induced loss of H3K27me3, to the N-terminus of the H3 gene coding sequence^4^. After transducing hNSCs with the lentivirus containing N-terminus tagged H3 constructs, or the untagged H3.3K27M, we compared H3K27me3 levels of our tagged and untagged hNSCs against the BT245 and SF7761 DMG lines using Western immunoblotting (Figure 1a). We observed no reduction in H3K27me3 levels in cells expressing either the N-terminus FLAG-HA tagged H3.1K27M (113.4% of untransduced NSC, p=0.467) or H3.3K27M (112.9% of untransduced NSC, p=0.530), despite observing that H3.3K27M expression was comparable between the tagged hNSC, untagged hNSC, and the DMG lines using an H3.3-specific anti-H3K27M antibody. These findings suggest that the addition of the N-terminal FLAG-HA tag may perturb the effect of H3K27M on suppressing total levels of H3K27me3.

This finding extended to other cell-types. Prior work evaluating the cellular hierarchy of diffuse midline gliomas has highlighted that tumors driven by H3.1K27M likely have a different cell-of-origin from those driven by H3.3K27M, with each arising from distinct waves of oligodendrocyte precursor cell (OPC) specification^16^. This raises the possibility that the histone mutations may exert their phenotypes distinctly in different cellular contexts. We therefore evaluated whether the lack of suppression of H3K27me3 observed in the setting of N-terminal tags on the mutant histone varied across different cell types. We tested 293T cells, human embryonic kidney cells that were used in the original demonstration^4^ of the dominant-negative effects of H3K27M on H3K27me3, and human embryonic stem cell (hESC)-derived OPCs transduced with the same N-terminus FLAG-HA tagged constructs. Expression of N-terminus tagged H3.3K27M did not reduce H3K27me3 levels in either 293Ts (Figure 1b, 120.8% of untransduced 293T, p=0.568) or OPCs (Figure 1c, 98.2% of untransduced OPC, p=0.914), though expression of N-terminal tagged H3.1K27M in HEK293Ts did mildly reduce H3K27me3 (45.6% of untransduced 293T, p=0.021).

To determine whether the tag peptide sequence itself interfered with the ability of H3K27M to suppress the H3K27me3 mark, we generated an additional panel of H3.3K27M expressing NSCs, in which the K27M mutants were tagged with either V5, FLAG or HA. Consistent with our observations with the FLAG-HA tag, we did not observe any suppression of the H3K27me3 marks with expression of any of the N-terminal tags on H3.3K27M (Figure 2a, 156.4%, 162.5%, 171.9%, and 227.4% of untransduced control for FLAG-HA, HA, FLAG, and V5 tagged samples respectively, p=0.296, 0.402, 0.286, and 0.058), suggesting that the presence of any N-terminal H3.3 tag may interfere with downstream functions of the K27M mutation.

**Figure 2.**
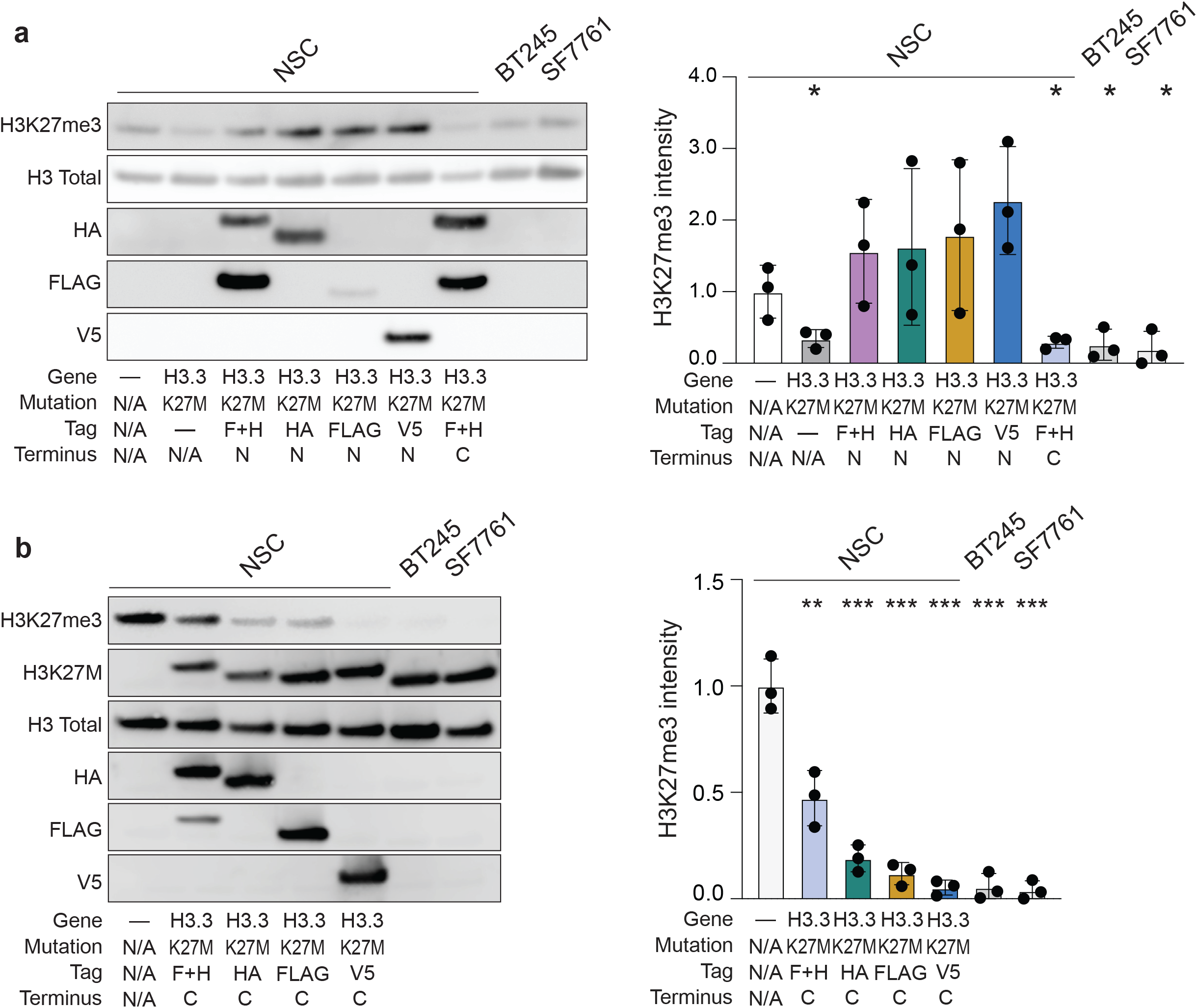
C-terminal tags do not interfere with H3K27M-mediated H3K27me3 loss. **a.** Left: Western immunoblot of acid-extracted lysates from hNSCs overexpressing H3.3K27M with the indicated N-terminal or C-terminal tag compared to the BT245 and SF7761 patient-derived DMG cell lines. Right: quantification of H3K27me3 : H3 ratios normalized to untransduced NSC control, n = 3 biological replicates. **b.** Left: Western immunoblot of acid-extracted lysates showing H3K27me3 levels in NSCs overexpressing H3.3K27M with the indicated C-terminal tags compared to BT245 and SF7761 cell lines. Right: quantification of H3K27me3 : H3 ratios normalized to untransduced NSC control, n = 3 biological replicates. Significance testing performed with two-sided unpaired t-test against untransduced NSC sample. Error bars, mean ± SEM of three replicates per condition. * = p<0.05, **=p<0.01, ***=p<0.001.

Next, we evaluated whether C-terminal tags similarly interfered with the ability of H3K27M to suppress H3K27me3 in NSCs. In contrast to N-terminal tags, the C-terminal FLAG-HA tagged H3K27M reduced H3K27me3 to a similar degree as its untagged K27M counterpart (Figure 2a, 29.4% of untransduced hNSCs, p=0.032). To determine whether the C-terminal FLAG-HA tag was unique in preserving the H3K27M-mediated reductions in H3K27me3, we added either a single FLAG, HA, or V5 tag, or FLAG-HA dual tag to the C-terminus of H3.3K27M. We found that NSCs expressing any of the C-terminally tagged H3K27M had significantly reduced H3K27me3 levels (Figure 2b, 19.0%, 11.9%, 5.3% of untransduced hNSCs for HA, FLAG, or V5 tagged models respectively, p<0.001 for all). Together, these results show that the addition of tags to the N-terminus of H3 nullify the H3K27M phenotype whereas C-terminal tags preserve it.

We next sought to understand the mechanism by which N-terminal tags drive the loss of H3K27M phenotype. In DMGs, direct binding of H3K27M to Polycomb Repressive Complex 2 (PRC2) is responsible for the reduced spreading of the H3K27me3 mark^7,17,18^. This suggests that either deficits in chromatin incorporation of H3K27M or altered binding dynamics of PRC2 at the N-terminal tail could be responsible for the changes observed in N-terminal tagged H3K27M. Though most residues associated with chaperone binding and nucleosome incorporation of H3 are found in its C-terminal core domain, we tested whether N-terminal epitope tagging may still inhibit efficient chromatin deposition of exogenous H3K27M in our models. To do this, we performed Cleavage Under Targets and Tagmentation (CUT&Tag) on hNSC lines expressing H3.3K27M with either the N-terminal or C-terminal FLAG-HA tag, H3.1K27M with an N-terminal FLAG-HA tag, or an empty vector control expressing HcRed. We found that H3.3K27M-FLAG signal coincided with active gene marker H3K4me2, especially at loci with a high density of variably expressed genes, such as the Hox gene cluster (Figure 3a). A broadly similar pattern was observed with the enrichment of endogenous H3.3K27M near gene bodies and transcriptionally active loci in the patient-derived DMG cell line DIPG-XIII (Figure 3a). Across all 14,946 parental hNSC H3K4me2 peaks, N-terminal tagged H3.3K27M showed a three-fold increase in mean anti-FLAG signal relative to N-terminal tagged H3.1K27M (3.21 vs 0.94 mean counts per million) and a near three-fold increase in fraction of reads in peaks (FRiP, 14.2% vs 4.9%), which had much more evenly distributed incorporation (Figure 3b, 3c). This pattern is consistent with the known transcription-associated incorporation pattern of endogenous H3.3 versus H3.1, which is incorporated throughout the genome during replication. We then compared relative deposition of N-terminal vs C-terminal tagged H3.3K27M and observed that both were robustly incorporated at H3K4me2 peaks, with 40% greater intensity in the N-terminally tagged samples for both anti-FLAG intensity and FRiP (3.21 vs 2.19 mean CPM, and 14.2% vs 10.2% FRiP, respectively). Together, these results indicate that both N- and C-terminally tagged exogenous H3K27M readily incorporate, and that retention of H3K27me3 in lines expressing N-terminal tagged H3K27M is likely due to altered interactions with chromatin writers at the H3 tail.

**Figure 3.**
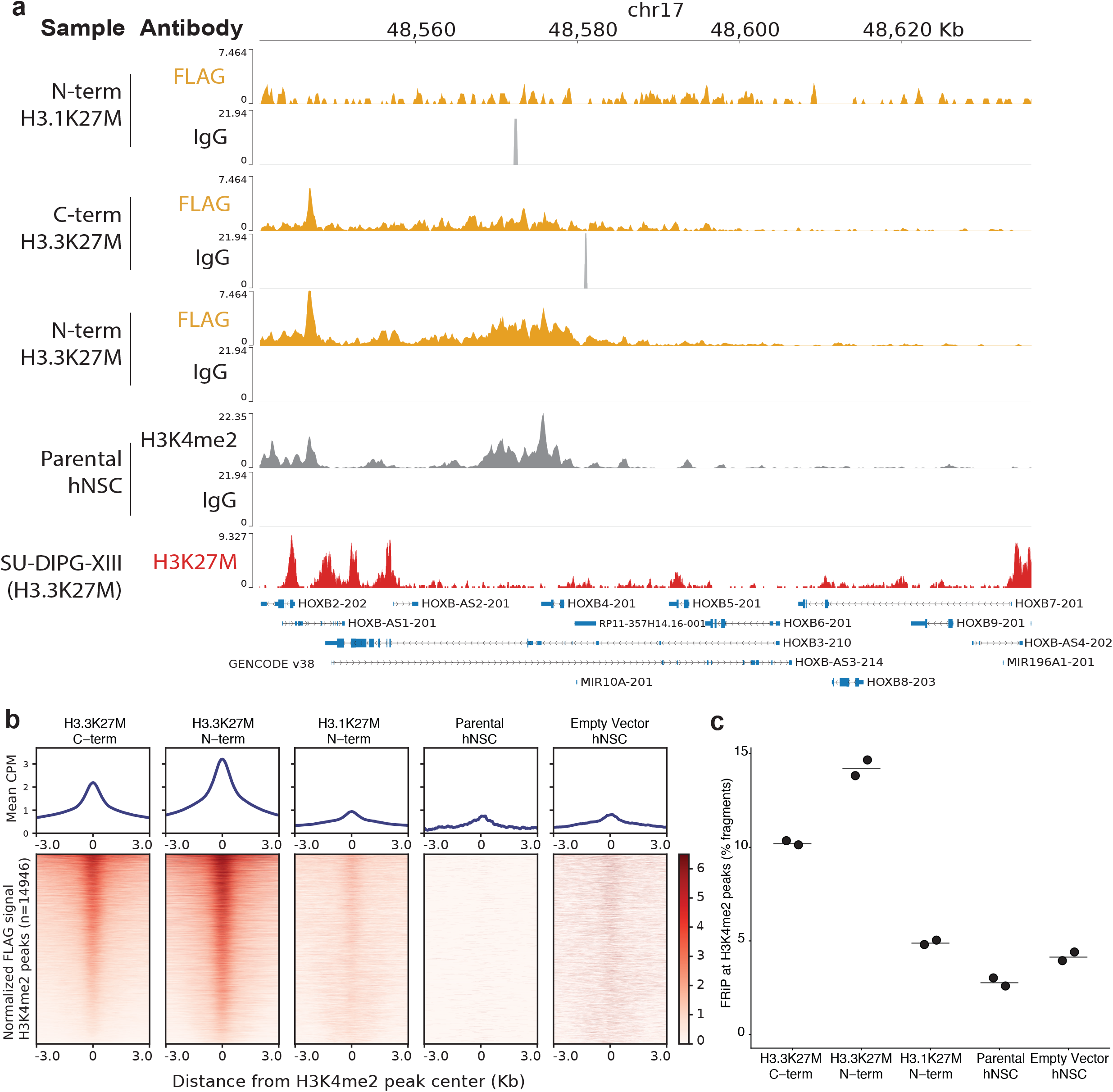
N-terminal and C-terminal tagged H3K27M are robustly incorporated into nucleosomes. **a.** Genome-browser tracks showing CUT&Tag signal from hNSC overexpressing N-terminal FLAG-HA tagged H3.1K27M, N- or C-terminal FLAG-HA tagged H3.3K27M, parental hNSC controls, or ChIP-seq signal for the SU-DIPG-XIII DMG cell line expressing endogenous H3.3K27M, from anti-FLAG, H3K27M, H3K4me2 antibodies or isotype controls as noted. Signal shown at representative HOX cluster locus. **b.** CUT&Tag signal at genome-wide H3K4me2 peaks: (Top) Metaplot of mean normalized anti-FLAG signal across all 14946 H3K4me2 peaks identified in control hNSCs, centered on peak midpoints and extending ± 3kb. (Bottom) Corresponding tornado plot showing normalized CUT&Tag signal at individual peaks over the same genomic interval. **c.** Fraction of total anti-FLAG reads located within H3K4me2 peaks (FRiP), across all 14946 H3K4me2 peaks identified in control hNSCs. n = 2 technical replicates

## Discussion

We found that adding N-terminal epitope tags to the mutant histone H3K27M attenuates its suppression of H3K27me3 across multiple cell types and distinct protein tags. Both N- and C-terminal tagged H3K27M incorporated into chromatin similarly, which indicates that N-terminal tags interfere with H3K27M-mediated H3K27me3 loss downstream of nucleosome formation. This presents the possibility that N-terminal tags disrupt H3 functions by altering interactions between H3 and chromatin readers or writers at the H3 tail (Figure 4). This has significant implications for the design and generation of models to study H3K27M mediated oncogenicity, particularly in the context of diffuse midline gliomas: N-terminal tags should be generally avoided, and rigorously validated if employed.

**Figure 4.**
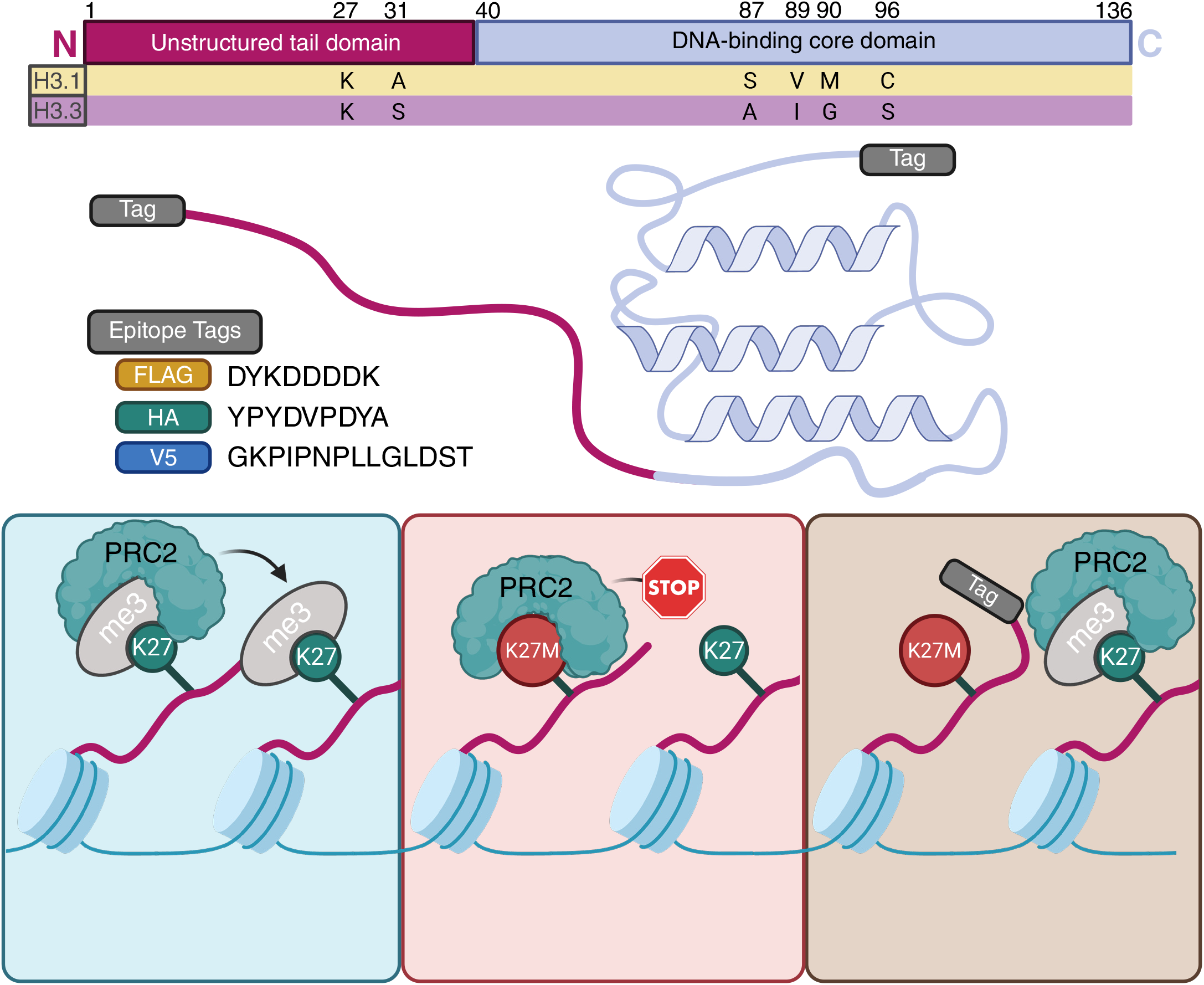
Graphical summary of findings: addition of N-terminal epitope tags interferes with PRC2 interactions with H3K27M.

These results suggest that portions of the N-terminal H3 tail distal to the DNA-binding core domain may have an important role in PRC2 binding. Cryo-EM structures of H3-bound PRC2 have shown that PRC2 makes multiple contacts to the H3 tail between residues R26 and K36^15,19,20^ proximal to the mutated K27M residue, whereas the residues on the N-terminal side of R26 typically remain unbound^17,21^. Alterations to these K27M-proximal residues, like H3K36 trimethylation, diminish K27M-dependent inhibition and PRC2 catalysis^15,22^. In contrast, less is known about how the distal N-terminal tail affects PRC2 function. There is evidence that such residues may still bind to PRC2: methylation of the H3K4 residue inhibits PRC2 activity through binding of an allosteric site on EED^15^. Additionally, truncation of the distal N-terminus of the histone tail reduced the PRC2 binding affinity and PRC2-inhibitory capacity of soluble H3K27M-tail-mimic peptides^23^. Our results are consistent with these reports, suggesting that preservation of the H3 N-terminus plays an important role in PRC2 activity on the H3 tail.

Though these results suggest that N-terminal tags alter the interaction between the PRC2 complex and the H3 tail, this study has several limitations. While we determine that tags disrupt the H3K27M phenotype downstream of chromatin deposition, this study does not demonstrate the precise mechanism behind this result. Additionally, we do not establish whether the same process that disrupts the loss of H3K27me3 in N-terminally tagged H3 would similarly affect other histone-binding proteins and PTMs at other sites on the H3 tail. Finally, our model systems rely on exogenous overexpression of the mutant histone which may not fully recapitulate the biology of endogenous histones. We have validated that our overexpression models have similar levels of H3K27M expression as DMG lines bearing the mutation at the endogenous locus, making it unlikely that this phenotype is due solely to supraphysiological expression of the mutation. Nonetheless, we cannot rule out that native control of the endogenous H3 loci may alter this phenotype in other models. It will thus be important to assess the effects of N- and C-terminal tags on function of the K27M mutations in the endogenous setting.

Our results support the idea that epitope tags should be used with caution and independently validated in each experimental model when studying histone biology. We found that appending any tag on the N-terminal tail nullifies H3K27M’s suppression of H3K27 trimethylation, making these models incompatible with the study of H3K27M-mediated oncogenesis in DMGs. Furthermore, these findings may explain the heterogeneity that we have observed in the magnitude of H3K27me3 reductions resulting from H3K27M expression across published results using different H3 tags. Taken together, these findings emphasize the need for careful validation of histone function when using modified histones and may warrant further investigation to determine whether N-terminal tags disrupt other H3-tail PTMs beyond H3K27me3.

## Materials and Methods

### Cell line models

#### Cell line sourcing and authentication

Human neural stem cells (hNSC) (ACS-5004) were purchased from American Type Culture Collection (Manassas, VA, USA). Lenti-X 293T cells were purchased from Takara Bio (Shiga, Japan). hESC-derived OPCs were purchased from EMD Millipore Sigma (SCR600). SF7761 patient-derived DMG cell line was purchased from Sigma Aldrich (SCC126). BT245 patient-derived DMG line (RRID:CVCL_IP13) was a gift from the Keith Ligon lab. Cells were routinely fingerprinted for identity validation and tested (at least every 3 months) for mycoplasma infection using the MycoAlert Mycoplasma Detection Kit (Lonza, LT07-318), according to the manufacturer’s instructions.

#### Culturing of hNSCs

hNSCs were maintained on adherent plates pretreated with Geltrex LDEV-Free Reduced Growth Factor Basement Membrane Matrix (Gibco, A1413201) for at least 30 minutes. hNSCs were grown in culture medium containing a 1:1 ratio of DMEM/F-12 (Invitrogen, 11330-032) and Neurobasal A (Invitrogen, 10888-022) with 1% each of HEPES Buffer Solution 1M (Thermo Fisher, 15630080), Sodium Pyruvate Solution 100 mM (Life Technologies, 11360070), MEM Non-Essential Amino Acids Solution 10 mM (Thermo Fisher, 11140050), Glutamax-I Supplement (Thermo Fisher, 35050061) and Penicillin/Streptomycin Solution (Life Technologies, 15140122). Media was supplemented with B-27 minus vitamin A (Invitrogen, 12587-010), epidermal growth factor (StemCell Tech. Inc., 78006), fibroblast growth factor (StemCell Tech. Inc., 78003) and 0.2% heparin solution (StemCell Tech. Inc., 07980). Culture media was replaced with fresh media every 48 hours and cells were dissociated using Accutase (StemCell Tech. Inc., 07922) to be passaged every 4 days.

#### Culturing of Lenti-X HEK293T cells

Lenti-X HEK293T were maintained on adherent plates and grown in culture medium of DMEM (Thermo Fisher, 11965092) supplemented with 10% of BenchMark Fetal Bovine Serum (GeminiBio, 100-106). Cells were dissociated using Trypsin-EDTA (0.25%) (Thermo Fisher, 25200114) to be passaged every 3 to 4 days.

#### Culturing of patient-derived DMG lines

BT245 and SF7761 cell lines were grown in tumor stem media containing a 1:1 ratio of DMEM/F-12 (Invitrogen, 11330-032) and Neurobasal-A (Invitrogen, 10888-022) with 1% each of HEPES Buffer Solution 1 M (Thermo Fisher, 15630080), sodium pyruvate solution 100 mM (Life Technologies, 11360070), MEM nonessential amino acids solution 10 mmol/L (Thermo Fisher, 11140050), Glutamax-I Supplement (Thermo Fisher, 35050061), and penicillin/streptomycin solution (Life Technologies, 15140122). Media was supplemented with B27 minus Vitamin A (Invitrogen, 12587-010), EGF, (StemCell Tech Inc., 78006), FGF (StemCell Tech Inc., 78003), and heparin solution 0.2% (StemCell Tech Inc., 07980), and PDGF-AA + PDGF-BB each 10ng/mL (Shenandoah Biotech 100-16 and 100-18). Cells were dissociated using Accutase (StemCell Tech. Inc., 07922) to be passaged every 4 days.

#### Culturing of OPCs

OPC cell line and optimized culture media kit was purchased from EMD Millipore Sigma and passaged according to manufacturer’s instructions. Cells were grown on Matrigel-coated adherent plates. Cells were dissociated using Accutase (StemCell Tech. Inc., 07922) to be passaged every 7-10 days.

### Lentiviral production and transductions

#### Plasmid generation

Primers were designed and ordered from Integrated DNA Technologies (Newark, NJ) for PCR amplification of desired inserts of H3F3A wild-type (WT) and H3F3A K27M with or without N or C terminal tags. PCR amplification was performed using full-length WT or K27M variations of the gene with N-terminal FLAG+HA tag as templates with the following cycle: 94°C for 2 seconds, followed by 30 cycles of 94°C for 10 seconds, 55 °C for 10 seconds, and 70°C for 15 seconds, with a final extension of 2 min at 70 °C. Insert DNA fragments were then purified using the QIAquick PCR Purification Kit (Qiagen, 28104) according to manufacturer’s instructions. Closed pLX312 vector obtained from the Broad Genetics Perturbation Platform (GPP) and insert DNA fragments were digested in a reaction mix consisting of 2 µL ClaI (New England BioLabs, R0197S), 1 µL EcoRV-HF (New England BioLabs, R3195S), 5 µL 10X NEBuffer (New England BioLabs, B6002S), 1 µg DNA, and ddH2O up to a total reaction mixture of 50 µL. Digestion proceeded for 1 hour at 37C. Digested insert DNA fragments were then purified using the QIAquick PCR Purification Kit (Qiagen, 28104) and digested vector was gel extracted using the QIAquick Gel Extraction Kit (Qiagen, 28704) according to manufacturer’s instructions. Digested insert DNA fragments were ligated to the open vector at a 3:1 molar ratio using 2 µL 10X Ligase Buffer (New England Biolabs, B0202S), 1 µL T4 Ligase (New England Biolabs, M0202L), and ddH2O up to a total reaction mixture of 20 µL. Ligation proceeded at room temperature for 2 hours and was heat-inactivated at 65°C for 10 minutes. Then, 10 μL of the ligation reaction was added to 50 μL of One Shot Stbl3 Chemically Competent E. coli (Invitrogen, C737303) and placed on ice for 30 minutes. The cells were then heat shocked at 42°C for 30 seconds and incubated on ice for 5 minutes. SOC media (Invitrogen, 15544034) was subsequently added to cells to allow for recovery and then shaken for 1 hour at 37°C. Transformants were plated on LB agar plates containing 100 μg/mL of carbenicillin and grown at 37°C for no more than 18 hours. Minipreps were performed using the QIAprep Spin Miniprep Kit (Qiagen, 27104) and sequences were verified by Plasmidsaurus whole plasmid sequencing. Once sequences were verified, Maxipreps were performed using the ZymoPURE II Plasmid Maxiprep Kit (Zymo Research, D4202).

#### Virus production

Lenti-X 293T cells (Takara Bio, 632180) were transfected with 10 μg of lentiviral expression vectors with packaging plasmids encoding PSPAX2 and VSVG using Lipofectamine 3000 (Thermo Fisher, L3000075). Media was replaced 6 hours after transfection. Supernatant containing lentivirus was collected 24 hours later, concentrated using the Lenti X-Concentrator (Takara, 631231) according to the manufacturer’s instructions, and collected in DMEM (Thermo Fisher, 11965092).

#### Transductions

hNSCs and Lenti-X 293T cells were seeded at 1.3 x 10^5^ cells/mL in adherent 10 cm dishes (CELLTREAT, 229620) and infected using no-spin infection 24 hours later. Dishes for hNSCs were pretreated with Geltrex LDEV-Free Reduced Growth Factor Basement Membrane Matrix (Gibco, A1413201) for at least 30 minutes. Infected cells were subsequently flow-sorted for GFP-positive cells 24 hours after infection.

#### Fluorescence-activated cell sorting

Cells were passaged as per routine cell culture methods and gated based on GFP expression with non-GFP-expressing cells used as negative controls. Cells were subsequently flow sorted on a BD FACSAria III directly into 15mL tubes containing appropriate media.

### Histone extraction and immunoblotting

Two-step histone extraction was performed on cells using the Histone Extraction Kit (Active Motif, 40028) according to manufacturer’s instructions. Supernatant was quantified with the Pierce 660 nm Protein Assay (Thermo Fisher, 22660), mixed with NuPAGE LDS Sample Buffer 4X (Invitrogen, NP0008), and heated at 90°C for 5 min. Equal amounts of protein for each sample were loaded alongside Cytiva Rainbow Molecular Weight Marker (Thermo Fisher, 45-001-591) and separated by SDS-PAGE on NuPAGE™ Bis-Tris Mini Protein Gels, 4–12% (Invitrogen, NP0321). Blots were subsequently dry transferred using the iBlot 2 Gel Transfer Device (Thermo Fisher, IB21001), placed in AdvanBlock-Chemi blocking solution (Advansta, R-03726-E10), and gently agitated for 1 hour at room temperature. Blots were then put in primary antibody and gently agitated at 4 °C overnight before being washed with 1X TBST three times for 15 minutes each. Antibodies against the following proteins were used at a 1:1000 dilution in this study: Histone H3-HRP (Cell Signaling, 12648), Tri-Methyl-Histone H3 (Lys27) (Cell Signaling, 9733), HA-Tag (Cell Signaling, 3724), V5-Tag (Cell Signaling, 13202), and DYKDDDDK Tag (Cell Signaling, 2368). Blots were then put in secondary antibody (Anti-rabbit IgG, HRP-linked antibody, Cell Signaling, 7074) (1:10,000 dilution) and gently agitated for one hour at room temperature before being washed with 1X TBST three times for 15 minutes each. Blots were saturated in SuperSignal West Femto Maximum Sensitivity Substrate (Thermo Fisher, 34095) and visualized on the Licor Odyssey M imaging system .

#### Immunoblot quantification

Quantification of the relative band intensities in the Western immunoblots was performed using the ImageJ/Fiji v2.14.0 software. Images with no overexposed pixels were selected for analysis. To measure band intensity, the Fiji Gel Analyzer tool was used to integrate pixel values across the length of the band. Background subtraction was performed on a per-lane basis to account for possible differences in exposure across the blot. To compare H3K27me3 intensity across samples, the ratio of H3K27me3 integrated intensity was normalized to H3-Total loading intensity. For samples with N≥3 replicates, statistical testing was performed with two-sided t-testing, assuming equal variance across samples, using Graphpad Prism to compare the H3K27me3:H3-Total ratio of the given sample against the H3K27me3:H3-Total ratio of the respective untransduced control line. Graphs of H3K27me3 levels were generated by normalizing the sample H3K27me3:H3-Total ratio against H3K27me3-H3Total ratio of the untransduced control line.

### CUT&Tag Sequencing

#### Data processing and analysis

Raw sequencing data were processed through the nf-core cutandrun pipeline (GRCh38/hg38 reference genome). Target reads were deduplicated, mitochondrial reads were removed, and per-replicate signal tracks were normalized to counts per million mapped reads (CPM). Replicate CPM bigWig files were averaged for each construct and antibody combination using deepTools bigwigAverage.

H3K4me2 peaks were called per replicate from deduplicated BAM files using MACS2 (paired-end BAMPE mode; q-value threshold 0.05; --nolambda --keep-dup all; effective genome size 2.86×10⁹ bp). A high-confidence H3K4me2 reference peak set was constructed by intersecting consensus peaks from the parental and HcRed EV controls.

Merged CPM bigWigs were scored across the robust H3K4me2 peak reference set using deepTools computeMatrix (reference-point mode; anchor at peak midpoint; ±3 kb flanking window; 25 bp bins; -- skipZeros). A single FLAG antibody sub-matrix was assembled including H3.3K27M-Cterm-FLAG-HA, H3.3K27M-Nterm-FLAG-HA, H3.1-K27M-Nterm-FLAG-HA, parental hNSC FLAG, and HcRed EV FLAG.

#### Heatmaps at H3K4me2 peaks

Tornado heatmaps were produced with deepTools plotHeatmap from the FLAG sub-matrix. Peaks were sorted in descending order by mean signal across the matrix. Color map: Reds; color scale upper bound set automatically to the 99th percentile of positive signal values.

#### Genome Browser Track Plots

Genome browser tracks were rendered with pyGenomeTracks from merged CPM bigWig files. The HOXB cluster locus was visualized at chr17:48,540,745–48,635,989, displaying FLAG-tag signal, H3K4me2 reference signal (Parental), and matched IgG negative controls for each of H3.1-K27M, H3.3-K27M, and parental cells, as well as published DIPG patient H3K27M ChIP-seq (GSE118099; SU-DIPG-XIII). Track y-axes were scaled to a shared per-antibody maximum computed from the 700-bin mean signal across the displayed locus (--scale-by-ab). Controls (Parental, HcRed EV) were auto-scaled independently to avoid compression of experimental tracks. A GENCODE v38 gene annotation track (merged transcripts, UCSC style) was appended below the signal tracks.

#### Fraction of Reads in Peaks (FRiP) analysis

To measure how much FLAG signal fell within H3K4me2 peaks, we calculated the fraction of fragment reads in peaks (FRiP) for each FLAG CUT&Tag library. For the numerator, we counted fragments overlapping control H3K4me2 peaks (14,946 regions) in each library with deepTools multiBamSummary (BED-file mode, --extendReads –samFlagInclude 64). These within-peak fragment counts were then divided by total mapped fragments per library, calculated by taking half the mapped reads reported by samtools idxstats. FRiP was reported as percentage for each biological replicate (n = 2 per construct), with bars marking the mean of the two replicates.

#### Restriction enzyme cloning oligos

H3.3-Tagless-Forward GGTGGTATCGATATGGCTCGTACAAAGCAGACTGCC

H3.3-Tagless-Reverse GGTGGTGATATCTCAAGCACGTTCTCCACGTATGCG

H3.3-Nterm-FLAG-Forward GGTGGTATCGATATGTACAAGGACGACGATGACAAGGCTCGTACAAAGCAGACTGCC

H3.3-Nterm-HA-Forward GGTGGTATCGATATGTACCCATACGATGTTCCAGATTACGCTGCTCGTACAAAGCAGACTGCC

H3.3-Nterm-V5-Forward GGTGGTATCGATATGGGTAAGCCTATCCCTAACCCTCTCCTCGGTCTCGATTCTACGGCTCGTACA AAGCAGACTGCC

H3.3-Nterm-HAorFLAGorV5-Reverse GGTGGTGATATCTCAAGCACGTTCTCCACGTATGCG

H3.3-Cterm-HAorFLAGorV5-Forward GGTGGTATCGATATGGCTCGTACAAAGCAGACTGCC

H3.3-Cterm-FLAG+HA-Reverse GGTGGTGATATCCTAAGCGTAATCTGGAACATCGTATGGGTACTTGTCATCGTCGTCCTTGTAAGC ACGTTCTCCACGTATGCG

## Acknowledgements and Funding Sources

NP was supported by a NIH NIGMS Medical Scientist Training Program Award (T32GM144273), a T32 Award (T32GM145407), and a NIH F30 predoctoral fellowship (F30CA298742). PB was supported by Alex’s Lemonade Stand Foundation, NIH R37 5R37CA255245-07, ChadTough Defeat DIPG Foundation, Prayers for Maria Foundation, Pediatric Brain Tumor Foundation, Giving for Gabby Fund, the McKenna Claire Foundation and We Love You Connie Foundation.

The content is solely the responsibility of the authors and does not necessarily represent the official views of the National Institute of General Medical Sciences or the National Institutes of Health.

## Contributions

NP and PB conceived of and designed the experiments. NP and GEB performed experiments, and along with JS and PB, interpreted results and wrote the manuscript. DFZ, JC, AFK, and AS assisted with experimental execution and interpretation. IG and OEF assisted with experimental execution.

